# ssRNA/DNA-Sensors via Embedded Strand-Displacement Programs in CRISPR/Cas9 Guides

**DOI:** 10.1101/264424

**Authors:** Noah Jakimo, Pranam Chatterjee, Joseph M Jacobson

**Author notes:** **Correspondence:** Correspondence and requests for materials should be addressed to N.J.

## Abstract

RNA and DNA profiles can help classify a variety of biological states, including disease, metabolism and cell type. In a proof-of-concept study on novel genetically encodable components for detecting single-stranded oligonucleotides, we engineered switchable CRISPR guide RNA (swigRNA) with Cas9 affinity that is conditional on sensing an oligo trigger molecule (trigRNA or trigDNA). RNA- and DNA-sensing swigRNAs that serve as off-to-on and on-to-off switches for Cas9 cleavage were achieved by computational design of hybridization dependencies in the guide. This study highlights functional swigRNA structures that implement toehold-gated strand-displacement with their triggers, and should merit further engineering and kinetic characterization.

---

Native ribonucleoprotein nucleases of CRISPR systems protect bacterial populations from viral proliferation by guide RNA (gRNA) that help to recognize and damage invading genomic material ^1^, and in some cases, result in collateral digestion of single-stranded RNA or DNA ^2–4^. While these CRISPR components confer certain sensing and actuation functionality, applications that demand more complex responses to physical and biochemical stimuli have led to an abundance of CRISPR-based engineering. To date, Cas9 from the Type II CRISPR system of *S. pyogenes* has dominated the focus of these efforts ^5–7^. Even when only considering modifications to the SpCas9 guide, CRISPR activity has been coupled to a variety of stimuli, such as light absorption ^8^ and emission ^9^, aptamer-associated small-molecules ^10^, RNA-binding proteins ^11^, ssDNA anti-sense oligos ^12^, and bacterial transcriptional regulators ^13^. In this work we demonstrate the first gRNA-embedded molecular programs for conditional assembly of guide and Cas9 in response to RNA triggers.

## Computational Design of Switchable Guide RNA

We implemented the two low and high Cas9-affinity states of swigRNA with a toehold-gated strand-displacement reaction. Several groups have previously employed this method for controlling transcription termination ^13^, RNA processing ^14^, translation initiation ^15^, aptamer fluorescence^16^, and nucleic acid-based logic ^17, 18^. Figure 1A illustrates its relation to the stages of swigRNA-Cas9 activity, which are listed as follows: 1) Insertions or extensions in the swigRNA molecule disrupt structural features essential for its complexing with Cas9. 2) The added sequence content includes an unpaired “toe-hold” domain to promote hybridization with the triggering oligonucleotide. 3) A sufficiently matched trigger that base-pairs with the swigRNA toehold proceeds to incrementally displace base-pairing within the swigRNA. 4) Intramolecular and intermolecular strand exchanges promote alternative base-pairings within the swigRNA. The resulting structure then rescues Cas9 affinity.

**Figure 1:**
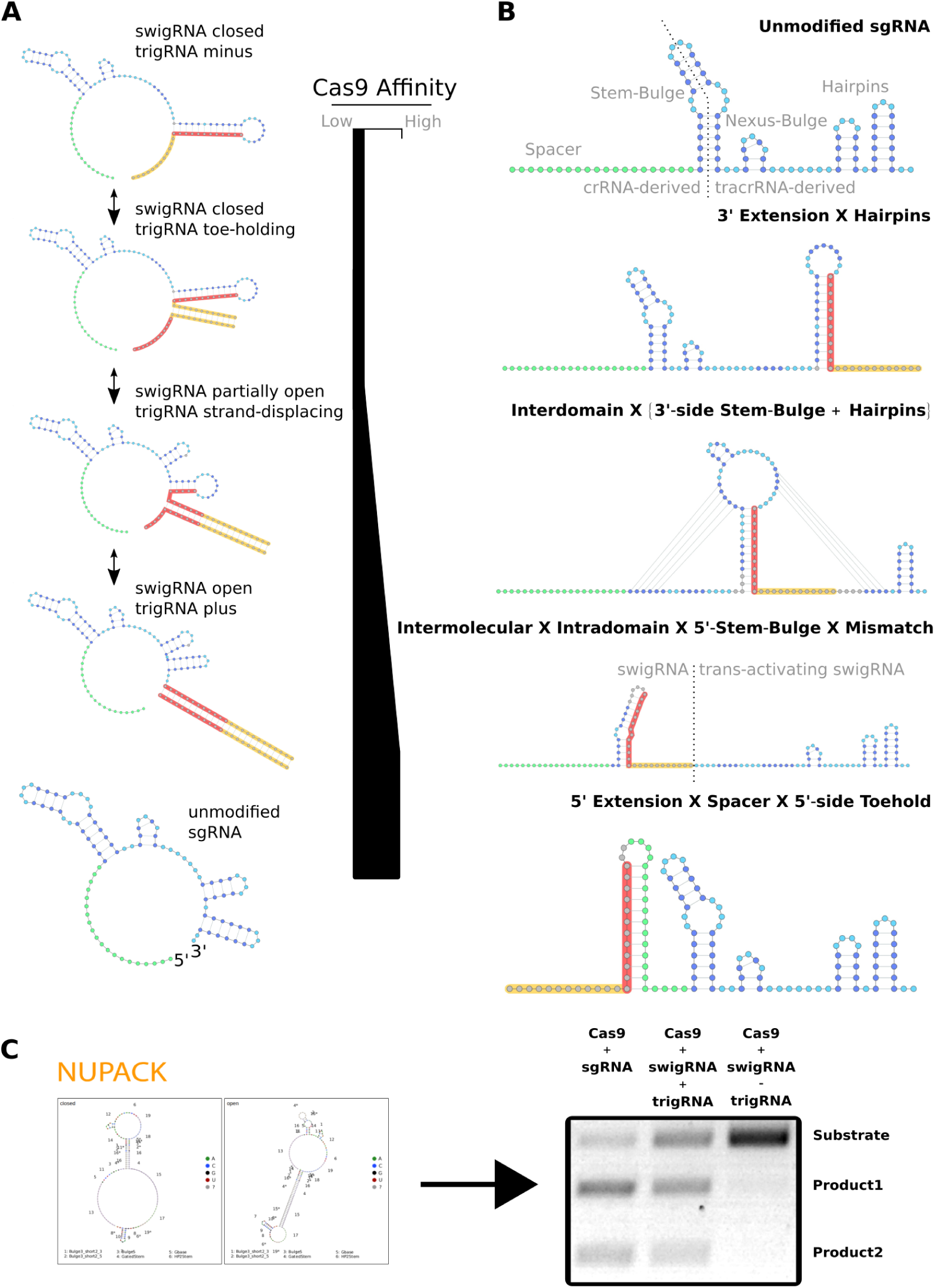
**A)** Illustration of the toehold-gated strand-displacement steps for switching swigRNA-Cas9 activity using trigger trigRNA. Colored domains denote target-defining spacer sequence (green), regions that map to positions in sgRNA that base-pair (dark blue), regions that map to positions in sgRNA that do not base-pair (light blue), and inserted swigRNA-unique sequences (gray). Highlighted domains denote domains in swigRNA and trigRNA that duplex based on toeholding (yellow) or strand-displacement (red). **B)** Modular strategies that impose RNA secondary structure constraints for swigRNA. **C)** Design, build, and test workflow for swigRNA evaluation.

We lay out general design principles for swigRNA (Figure 1B) by incorporating other studies on the effectiveness of gRNA variants that contain target-independent sequence and structural changes ^19, 20^. These principles exploit domains of a guide that can tolerate insertions or structure-preserving sequence substitutions. They allow terminal extensions, as well as resizing loops and stems in either the 3’-proximal hairpins or at the native duplex junction between CRISPR RNA (crRNA) and trans-activating CRISPR RNA (tracrRNA). We generated sequences that satisfy our design constraints using the nucleic acid computational design and analysis package, NUPACK ^21^. We were able to specify stable switching behavior by including OFF- and ON-state models in co-optimization tasks. In Figure 1C, we demonstrate functional switching by a standout design, which was evaluated by an *in vitro* Cas9 cleavage and gel shift assay ^22^.

## Validation of the Toehold-Gated Strand-Displacement Mechanism

We further assessed the same design (Figure 2) to validate the role of toehold-gated strand-displacement. As shown in Figure 2A, in the absence of trigRNA, only a marginal fraction of target DNA substrate was cleaved after 8 hours of digestion with Cas9 and swigRNA. By contrast, a time course that included a two-fold excess of trigRNA yielded cleavage on roughly half of the targets within the first hour. In support of our proposed mechanism, we found neither toeholding nor strand-displacing domain was sufficient for a trigger to induce cleavage. (Figure 2B and 2C – last lanes).

**Figure 2:**
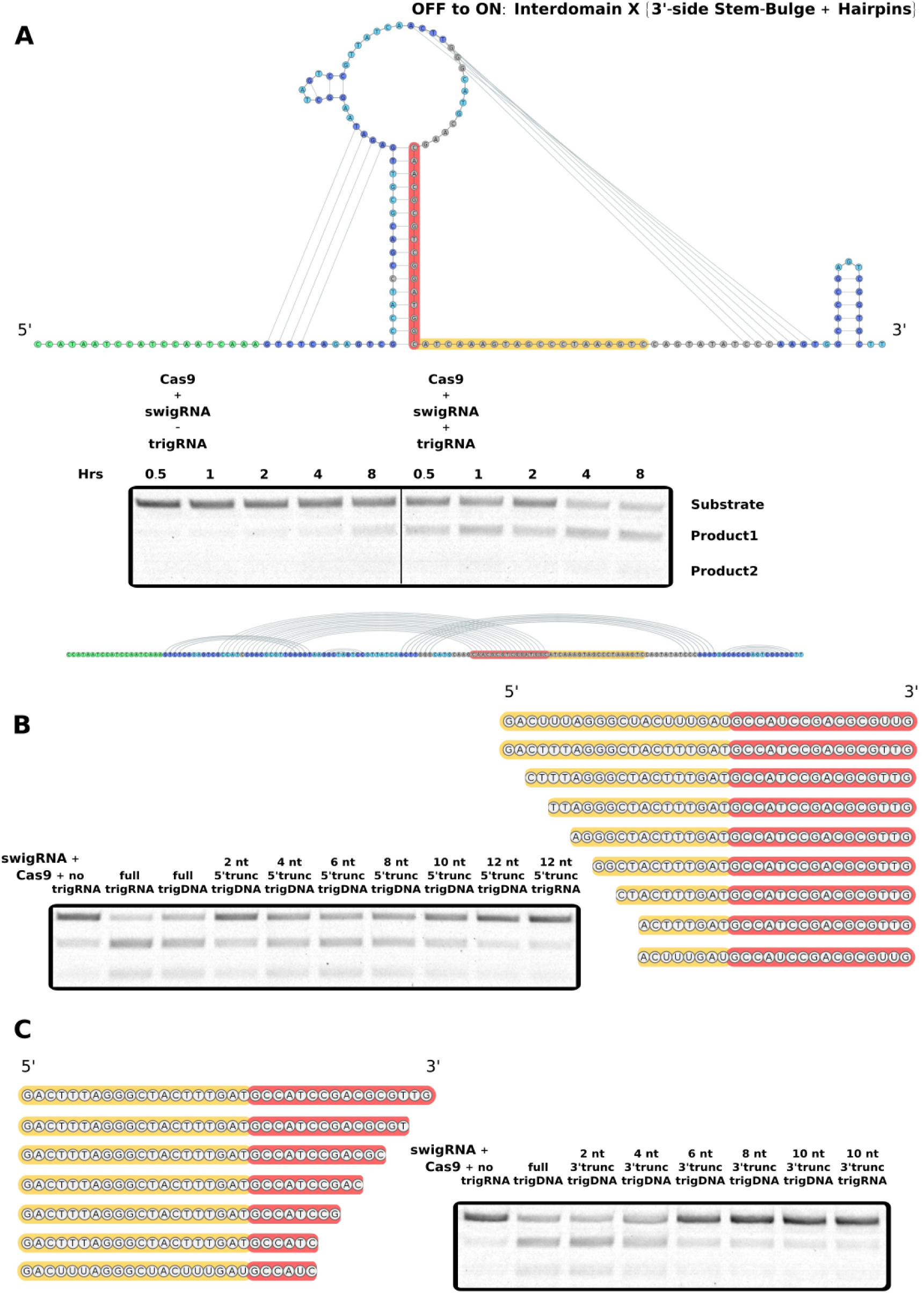
**A)** 2D and 1D representations of base-pairing in a swigRNA that is tested over an 8 hour time-course with and without trigRNA. **B)** Cas9 digest reactions run with 5’-end truncations of trigRNA or trigDNA. **C)** Cas9 digest reactions run with 3’-end truncations of trigRNA or trigDNA.

Like trigRNA, full-length trigDNA was also effective at inducing switching, yet with slightly lower efficiency. We expect this decrease in performance results from generally higher thermal stability of RNA:RNA hybridization relative to RNA:DNA. Nonetheless, we tested a variety of trigDNA with partial terminal truncations in order to probe the length of toeholding and strand-displacement needed for triggering swigRNA (Figure 2B and 2C). We observed 5’-end truncations in the toeholding domain of trigger gradually reduced digestion rates. While 2–4 nt 3’-end truncations in the strand-displacing domain had minimal effects on cleavage yields, those that extended beyond 5 nt showed drastic reduction.

We take these results to mean that toehold domain binding alone was insufficient to promote strand exchanges for stronger swigRNA-Cas9 affinity. Toeholding, however, facilitated invasion and displacement of base-pairing that inhibited Cas9 binding. Since near-complete displacement enabled formation of active swigRNA-Cas9, this switching after only partial invasion could suggest the remaining displacement occurred by thermal dehybridization (“breathing”), was promoted by intermediate strand-exchange structures, and was then stabilized by Cas9 binding.

## Demonstration of OFF-to-ON and ON-to-OFF Switchable Guide RNA

We confirm the same design strategy generates additional swigRNA that have off-to-on performance for other triggers and targets (Figure 3A). We were surprised to discover alternative design strategies produced opposite on-to-off switching (Figure 3B). These on-to-off swigRNA embed the entire toehold-gated strand-displacement program – in different orientations – within the junction of crRNA and tracrRNA. We expect this region is sterically sensitive to trigger hybridization for the stem lengths used in these designs. Lastly, we validate more swigRNA hybridization strategies that produce our intended off-to-on switching (Figure 3C).

**Figure 3:**
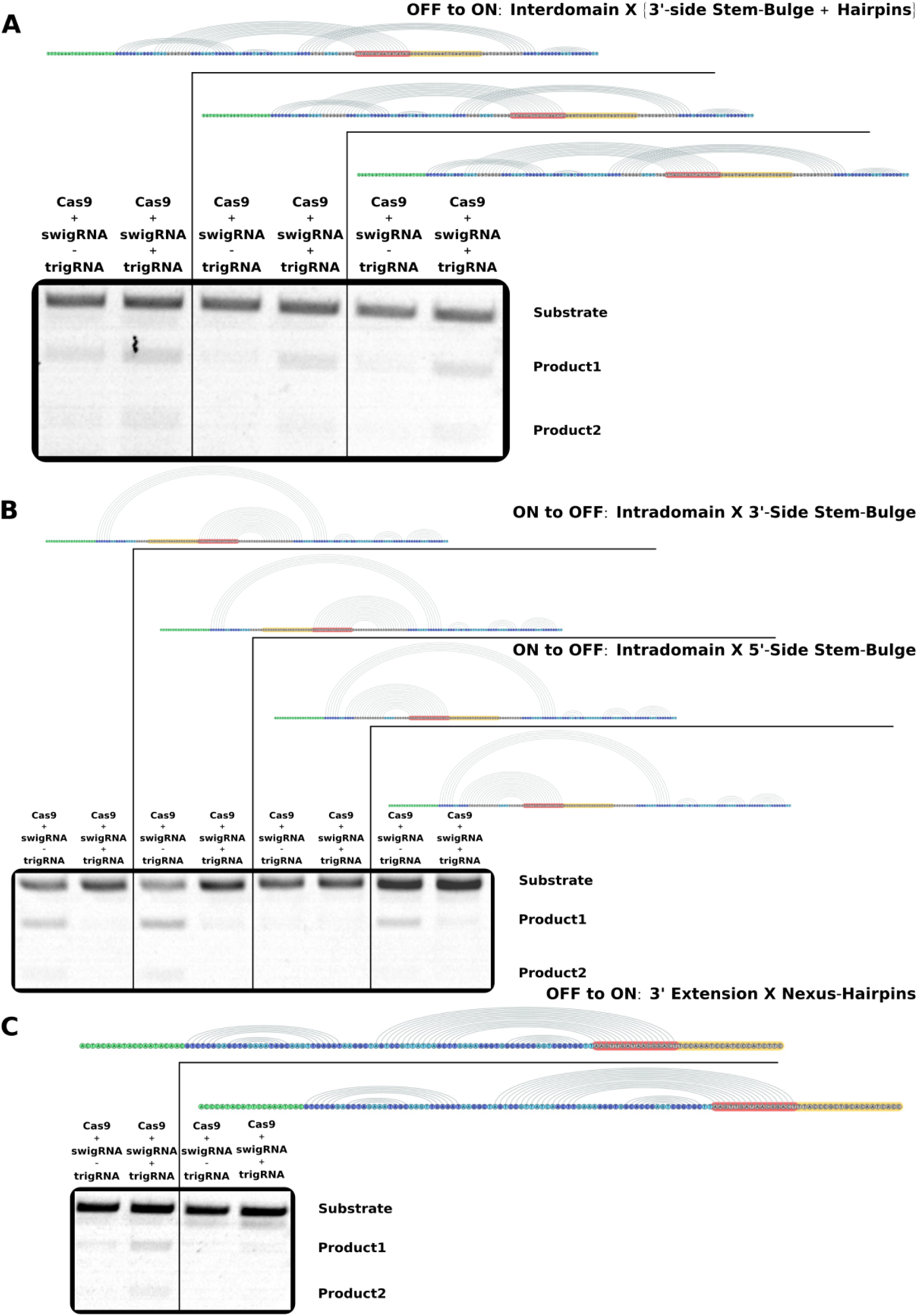
**A)** Evaluation of additional off-to-on swigRNA based on interdomain hybridization between the stem-bulge and hairpins. **B)** Demonstration of on-to-off swigRNA based on intradomain hybridization within the stem-bulge. **C)** Evaluation of alternative off-to-on swigRNA based on hybridization between a 3’ terminal extension and the nexus and hairpins.

Our work reveals several methods for coupling activity of the CRISPR/Cas9 system to sensing ssRNA or ssDNA molecules. We have also established a workflow that can now be applied to broader design libraries for new DNA targets and RNA triggers. We plan to correspond predicted and actual swigRNA structures using techniques, such as SHAPE-seq ^23^, in order to adjust our design principles for further generality.

## Methods

### *In vitro* Cas9 Cleavage Assays

IDT DNA Ultramers of reverse-complimented swigRNA designs were transcribed using the NEB HiScribe T7 Quick High Yield RNA Synthesis Kit and purified using the Qiagen RNeasy Mini kit. Each cleavage reaction consisted of 30 nM NEB Cas9, *S. pyogenes* nuclease, 30 nM swigRNA, 3 nM PCR amplified DNA target substrate, and 60 nM HPLC-purified synthetic trigRNA or trigDNA from IDT. Unless otherwise specified, digests incubated 4 hours before 2% TAE agarose gel electrophoresis.

### Nupack swigRNA Design

We used the online NUPACK nucleic acid sequence design server (http://nupack.org/design/new) to co-optimize active and inactive states of swigRNA. We wrote Python code to both parametrically generate an input NUPACK script and parse the output sequences into genbank format.

## Author Contributions

N.J. conceived and designed experiments. N.J. implemented computational workflows. N.J. and P.C. carried out experiments. N.J. wrote the paper. N.J., P.C., and J.M.J. reviewed the paper. J.M.J. supervised the project.

## Acknowledgments

We thank Dr. Neil Gershenfeld and Dr. Shuguang Zhang for shared lab equipment.

